# Maximum-entropy and representative samples of neuronal activity: a dilemma

**DOI:** 10.1101/329193

**Authors:** P.G.L. Porta Mana, V. Rostami, E. Torre, Y. Roudi

## Abstract

The present work shows that the maximum-entropy method can be applied to a sample of neuronal recordings along two different routes: (1) apply to the sample; or (2) apply to a larger, unsampled neuronal population from which the sample is drawn, and then marginalize to the sample. These two routes give inequivalent results. The second route can be further generalized to the case where the size of the larger population is unknown. Which route should be chosen? Some arguments are presented in favour of the second. This work also presents and discusses probability formulae that relate states of knowledge about a population and its samples, and that may be useful for sampling problems in neuroscience.

## 1 Introduction: maximum-entropy and recordings of neuronal activity

Suppose that we have recorded the firing activity of a hundred neurons, sampled from a particular brain area. What are we to do with such data? Gerstein, Perkel, Dayhoff [1] posed this question very tersely (our emphasis):

> The principal conceptual problems are (1) *defining cooperativity or functional grouping* among neurons and (2) *formulating quantitative criteria* for recognizing and characterizing such cooperativity.

These questions have a long history, of course; see for instance the 1966 review by Moore et al. [2]. The neuroscientific literature has offered several mathematical definitions of ‘cooperativity’ or ‘functional grouping’ and criteria to quantify it.

One such quantitative criterion relies on the maximum-entropy or relative-maximum-entropy method [3–7]. This criterion has been used in neuroscience at least since the 1990s, applied to data recorded from brain areas as diverse as retina and motor cortex [8–24], and it has been subjected to mathematical and conceptual scrutiny [25–30].

‘Cooperativity’ can be quantified and characterized with maximum-entropy methods in several ways. The simplest way roughly proceeds along the following steps. Consider the recorded activity of a sample of *n* neurons.

1. The activity of each neuron, a continuous signal, is divided into *T* time bins and binarized in intensity, and thus transformed into a sequence of digits ‘0’s (inactive) and ‘1’s [cf. 31; 32]. Let the variable *s_i_*(*t*) ∈ {0,1} denote the activity of the *i*th sampled neuron at time bin *t*. Collectively denote the *n* activities with *s*(*t*): =(*s*_1_(*t*),…, *s_n_*(*t*)). The population-averaged activity at that bin is 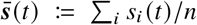. If we count the number of distinct pairs of active neurons at that bin we combinatorially find 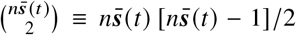. There can be at most 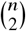 simultaneously active pairs, so the population-averaged pair activity is 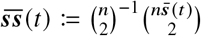. With some combinatorics we see that the population-averaged activity of *m*-tuples of neurons is

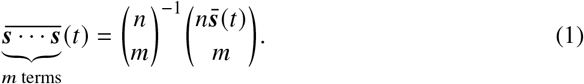 For brevity let us agree to simply call ‘activity’ the average 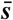, ‘pair-activity’ the average 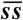, and so on.
2. Construct a sequence of relative-maximum-entropy distributions for the activity 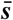, using this sequence of constraints:

- the time average of the activity: 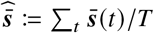;
- the time averages of the activity and of the pair-activity 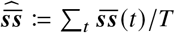;
- …
- the time averages of the activity, of the pair-activity, and so on, up to the *k*-activity. Call the resulting distributions 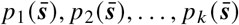. The time-bin dependence is now absent because these distributions can be interpreted as referring to any one of the time bins *t*, or to a new time bin (in the future or in the past) containing new data. We also have the empirical frequency distribution of the total activity, 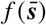, counted from the time bins.
3. Now compare the distributions above with one another and with the frequency distribution, using some probability-space distance like the relative entropy or discrimination information [33; 4; 34; 5]. If we find, say, that such distance is very high between *p*_1_ and *f*, very low between *p*_2_ and *f*, and is more or less the same between all *p_m_* and *f* for *m* ⩾ 2, then we can say that there is a ‘pairwise cooperativity’, and that any higher-order cooperativity is just a reflection or consequence of the pairwise one. The reason is that the information from higher-order simultaneous activities did not lead to appreciable changes in the distribution obtained from pair activities.

The protocol above needs to be made precise by specifying various parameters, such as the width of the time bins or the probability distance used.

We hurry to say that the description just given is just *one* way to quantify and characterize cooperativity and functional grouping, not *the only* way. It can surely be criticized from many points of view. Yet, it is quantitative and bears a more precise meaning than an undefined, vague notion of ‘cooperativity’. Two persons who apply this procedure to the same data will obtain the same numbers. Different protocols can be based on the maximum-entropy method, for instance protocols that take into account the activities or pair activities of specific neurons rather than population averages, or even protocols that take into account time dependence.

The purpose of the present work, based on the discussion and results of [35], is not to assess the merits of maximum-entropy methods with respect to other methods. Its main purpose is to show that there is a problem in the particular application, sketched above, of the maximum-entropy method to the activity of the recorded neurons. We believe that this problem and possible misapplication is at the root of some observations made in the literature [27]. This problem also extends to more complex applications of the method, possibly excepting versions that use ‘hidden’ neurons [36–39].

The problem is that the recorded neurons are a *sample* from a larger, unrecorded population, but the maximum-entropy method as applied above is treating them as isolated from the rest of the brain. Hence the results it provides cannot rightfully be extrapolated. We will give a mathematical proof of this. Let us first analyse this issue in more detail.

Suppose that the neurons were recorded with electrodes covering an area of some square millimetres [cf. 40]. This recording is a sample of the activity of the neuronal population under the recording device, a population that can amount to tens of thousands of neurons [41]. We could even consider the recorded neurons as a sample of a brain area more extended than the recording device.

The characterization of the cooperativity of the recorded sample would have little meaning if we did not expect its results to generalize to a larger, unrecorded population – at the very least the one under the recording device. In other words, we expect that the conclusions drawn with the maximum-entropy methods about the sampled neurons should somehow extrapolate to unrecorded neurons in some larger area, from which the recorded neurons were sampled. In statistical terms we are assuming that the recorded neurons are a *representative sample*^1^ of some larger neuronal population. Probability theory tells us how to make inferences from a sample to the larger population from which it is sampled (see references below).

We can apply the maximum-entropy method to the sample, as described in the above protocol, to generate probability distributions for the activity of the sample. But, given that our sample is representative of a larger population, we can also apply the maximum-entropy method to the larger (unrecorded) population. The constraints are the same: namely the time averages of the sampled data – in fact they constitute representative data about the larger population as well. The method thus yields a probability distribution for the larger population, and the distribution for the sample is then obtained by marginalization from that. The problem is that *the distributions obtained from these two applications differ*. Which choice is most meaningful?

In this work we develop the second way [sketched in 35 § 3] of applying the maximum-entropy method, at the level of the larger population, and show that its results differ from the application at the sample level. We also consider the case where the size of the larger population is unknown.

To apply the maximum-entropy method to the larger, unsampled population, it is necessary to use probability relations relevant to sampling [45; 46 parts I, VI; 47 ch. 3; 35]. The relations we present are well-known in survey sampling and in the pedagogic problem of drawing from an urn without replacement, yet they are somewhat hard to find explicitly written in the neuroscientific literature. We present and discuss them in the next section. A minor purpose of this paper is to make these relations more widely known, because they can be useful independently of maximum-entropy methods.

The notation and terminology in the present work follow ISO and ANSI standards [48–52] but for the use of the comma ‘,’ to denote logical conjunction. Probability notation follows Jaynes [47]. By ‘probability’ we mean a degree of belief which ‘would be agreed by all rational men if there were any rational men’ [53].

## 2 Probability relations between population and sample

We have already introduced the notation for the sample neurons. We introduce an analogous notation for the 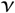 neurons constituting the larger population, but using the corresponding Greek letters: *σ_ι_*(*t*) is the activity of the *ι*th neuron at time bin *t*, 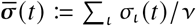 is the activity at that bin averaged over the larger population, and so on.

The probability relations between sample and larger population are valid at every time bin. As we mentioned above, the maximum-entropy distribution refers to any time bin or to a new bin. For these reasons we will now omit the time-bin argument ‘(t)’ from our expressions.

Probabilities refer to statements about the quantities we observe. We use the standard notation:

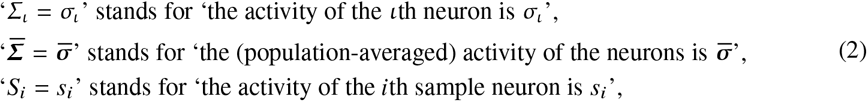

and similarly for other quantities.

If *K* denotes our state of knowledge – the evidence and assumptions backing our probability assignments – our uncertainty about the full activity of the larger population is expressed by the joint probability distribution

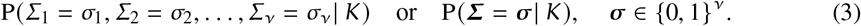

Our uncertainty about the state of the sample is likewise expressed by

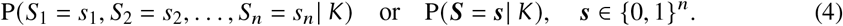

The theory of statistical sampling is covered in many excellent texts, for example Ghosh & Meeden [45] or Freedman, Pisani, & Purves [46 parts I, VI]; a summary can be found in Jaynes [47 ch. 3].

We need to make an initial probability assignment for the state of the full population before any experimental observations are made. This initial assignment will be modified by our experimental observations, and these can involve just a sample of the population. Our state of knowledge and initial probability assignment should reflect that samples are somehow representative of the whole population.

In this state of knowledge, denoted *I*, we know that the neurons in the population are biologically or functionally similar, for example in morphology or the kind of input or output they receive or give. But we are completely ignorant about the physical details of the individual neurons. Our ignorance is therefore symmetric under permutations of neuron identities. This ignorance is represented by a probability distribution that is symmetric under permutations of neuron identities; such a distribution is usually called *finitely exchangeable* [54; 45 ch. 1]. We stress that this probability assignment is just an expression of the symmetry of our *ignorance* about the state of the population, not an expression of some biologic or physical symmetry or identity of the neurons.

The *representation theorem for finite exchangeability* states that, in the state of knowledge I, the symmetric distribution for the full activity is completely determined by the distribution for its population-average:

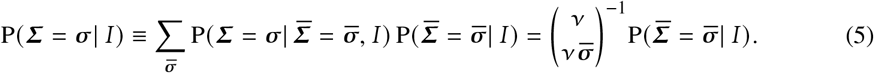

The equivalence on the left is just an application of the law of total probability; the equality on the right is the statement of the theorem. This result is intuitive: owing to symmetry, we must assign equal probabilities to all 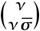 activity vectors with 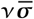 active neurons; the probability of each activity vector is therefore given by that of the average activity divided by the number of possible vector values. Proof of this theorem and generalizations to non-binary and continuum cases are given by de Finetti [55], Kendall [56], Ericson [57], Diaconis & Freedman [58; 59], Heath & Sudderth [60].

Our uncertainties about the full population and the sample are connected via the conditional probability

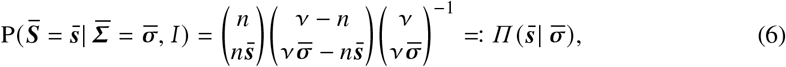

which is a hypergeometric distribution, typical of ‘drawing without replacement’ problems. The combinatorial proof of this expression is in fact the same as for this class of problems [47 ch. 3; 61 § 4.8.3; 62 § II.6].

Using the conditional probability above we obtain the probability for the activity of the sample:

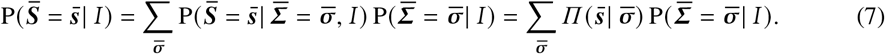

It should be proved that the probability distribution for the full activity of the sample is also symmetric and completely determined by the distribution of its population-averaged activity:

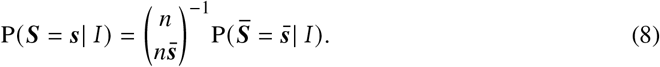

This is intuitively clear: our initial symmetric ignorance should also apply to the sample. The distribution for the sample (7) indeed satisfies the same representation theorem (5) as the distribution for the full population.

The conditional probability 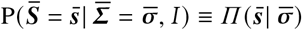, besides relating the distributions for the population and sample activities via marginalization, also allows us to express the expectation value of any function of the sample activity, 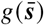, in terms of the distribution for the full population, as follows:

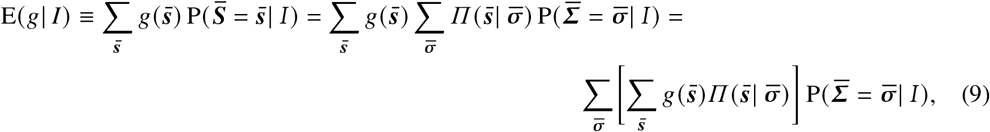

where the second step uses eq. (7). The last expression shows that the expectation of the function 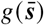 is equal to the expectation of the function 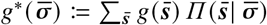.

The final expression in eq. (9) is important for our maximum-entropy application: the requirement that the function *g*, defined for the sample, have a value *c* obtained from observed data, *translates into a linear constraint for the distribution of the full population*:

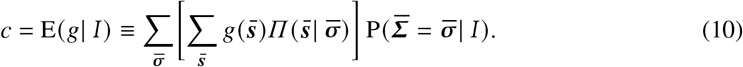

In particular, when the function *g* is the *m*-activity of the sample, 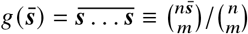, we find

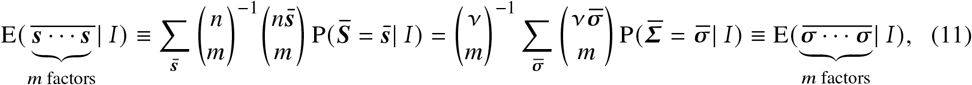

that is, *the expected values of the m-activities of the sample and of the full population are equal*. The proof of the middle equality uses the expression for the *m*th factorial moment of the hypergeometric distribution and can be found in Potts [63]. Similar relations can be found for the raw moments 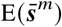 and 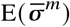, which can be written in terms of the product expectations using eq. (1).

Thus, in a maximum-entropy application, when we require the expectation of the *m*-activity of a sample to have a particular value, we are also requiring the expectation of the *m*-activity of the full population to have the same value.

These expectation equalities between sample and full population should not be surprising: we intuitively *expect* that the proportion of coloured balls sampled from an urn should be roughly equal to the proportion of coloured ball contained in the urn. The formulae in the present section formalize and mathematically express our intuition. The hypergeometric distribution 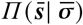 plays an important role in this formalization. A look at its plot, fig. 1, reveals that it is a sort of ‘fuzzy identity transformation’, or fuzzy Kronecker delta, between the 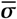-space {0,…, *ν*} and 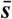-space {0,…, *n*}. From eq. (8) we thus have that

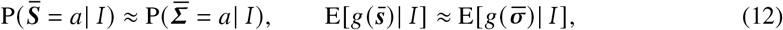

**Figure 1:**
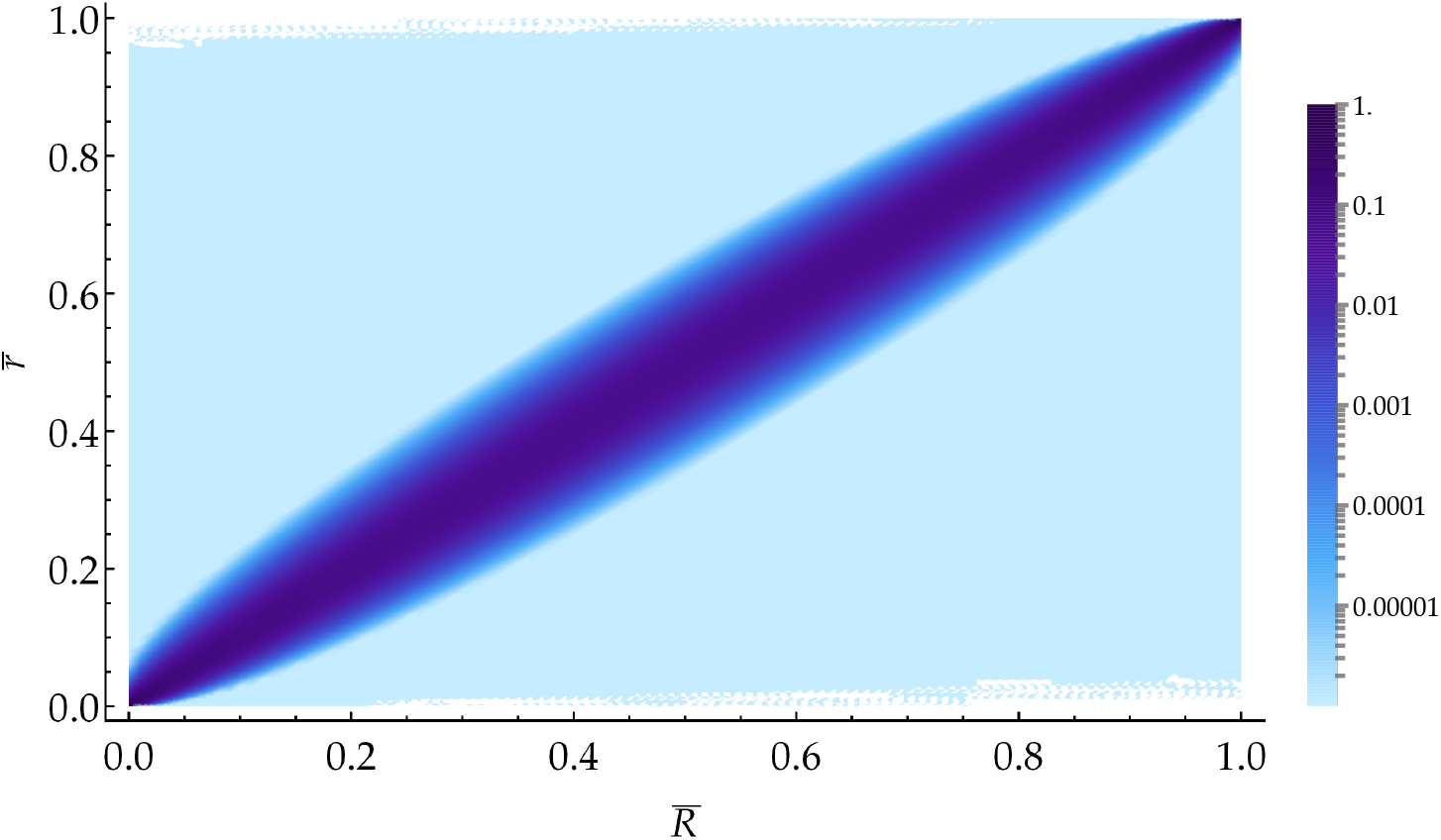
Log-density plot of the hypergeometric distribution 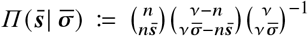 for *ν* = 5000, *n* = 200. (Band artifacts may appear in the colourbar depending on your PDF viewer.)

where *g* is any smooth function defined on [0,1]. These approximate equalities express the intuitive fact that *our uncertainty about the sample is representative of our uncertainty about the population and about other samples*, and vice versa. When 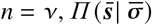, becomes the identity matrix and the approximate equalities above become exact – of course, since we have sampled the full population.

But the approximate equalities above may miss important features of the two probability distributions. In the next section we will in fact emphasize their differences. If the distribution for the population average 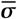 is bimodal, for example, the bimodality can be lost in the distribution for the sample average 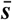, owing to the coarsening effect of 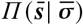.

## 3 Maximum-entropy: sample level vs full-population level

In the previous section we have seen that observations about a sample can be used as constraints on the distribution for the activity of the full population. Let us use such constraints with the maximumentropy method. Suppose that we want to constrain *m* functions of the sample activity, vectorially written ***g*** = (*g*_1_,…, *g_m_*), to *m* values ***c*** = (*c*_1_,…, *c_m_*). These functions are typically *k*-activities 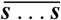, and the values are typically the time averages of the observed sample, as discussed in § 1: 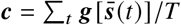.

Let us apply the relative-maximum-entropy method [6; 7] directly to sampled neurons; denote this approach by *I*_s_. Then we apply the method to the full population of neurons, most of which are unsampled; denote this approach by *I*_p_.

Applied directly to the sampled neurons, the method yields the distribution

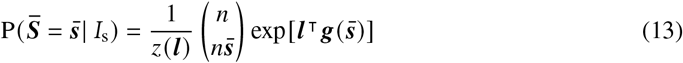

where *z*(***l***) is a normalization constant. The binomial in front of the exponential appears because we must account for the multiplicity by which the population-average activity 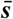 can be realized: 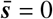 can be realized in only one way (all neurons inactive), 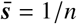 can be realized in *n* ways (one active neuron out of *n*), and so on. This term is analogous to the ‘density of states’ in front of the Boltzmann factor in statistical mechanics [64 ch. 16]. The *m* Lagrange multipliers ***l*** = (*l*_1_,…, *l_m_*) must satisfy the *m* constraint equations

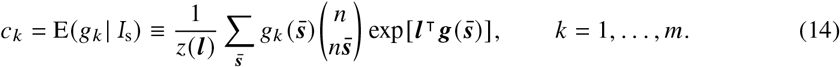

Applied to the full population, using the constraint expression (10) derived in the previous section, the method yields the distribution for the full-population activity

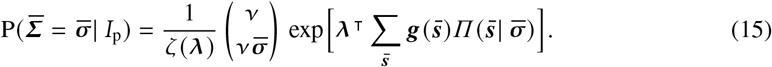

The *m* Lagrange multipliers ***λ*** = (*λ*_1_,…, *λ_m_*) must satisfy the *m* constraint equations

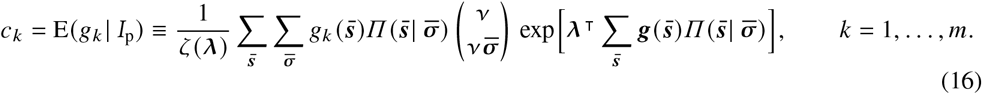

We obtain the distribution for the sample activity by marginalization, using eq. (8):

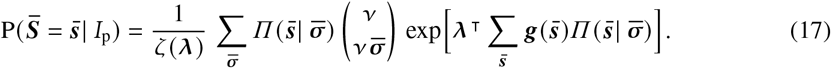

The distributions for the sample activity, eqs (17) and (13), obtained with the two approaches *I*_s_ and *I*_p_, are different. From the discussion in the previous section we expect them to be vaguely similar; yet they cannot be exactly equal, because their equality would require the 2*m* quantities ***λ*** and ***l*** to satisfy the constraint equations (16) and (14), and in addition also the *n* equations 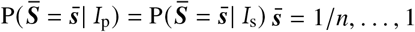 (one equation is taken care of by the normalization of the distributions). We would have a set of 2*m* + *n* equations in 2*m* unknowns.

Hence, *the applications of maximum-entropy at the sample level and at the full-population level are inequivalent*. They lead to numerically different distributions for the sample activity ***s***.

The distribution obtained at the sample level will show different features from the one obtained at the population level, like displaced or additional modes or particular tail behaviour. We show an example of this discrepancy in fig. 2, for *ν* = 10 000, *n* = 200, and the two constraints

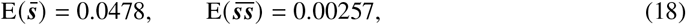

which come from the actual recording of circa 200 neurons from macaque motor cortex [30]. The distribution obtained at the population level (blue triangles) has a higher and displaced mode and a quite different behaviour for activities around 0.5 than the distribution obtained at the sample level (red squares).

**Figure 2:**
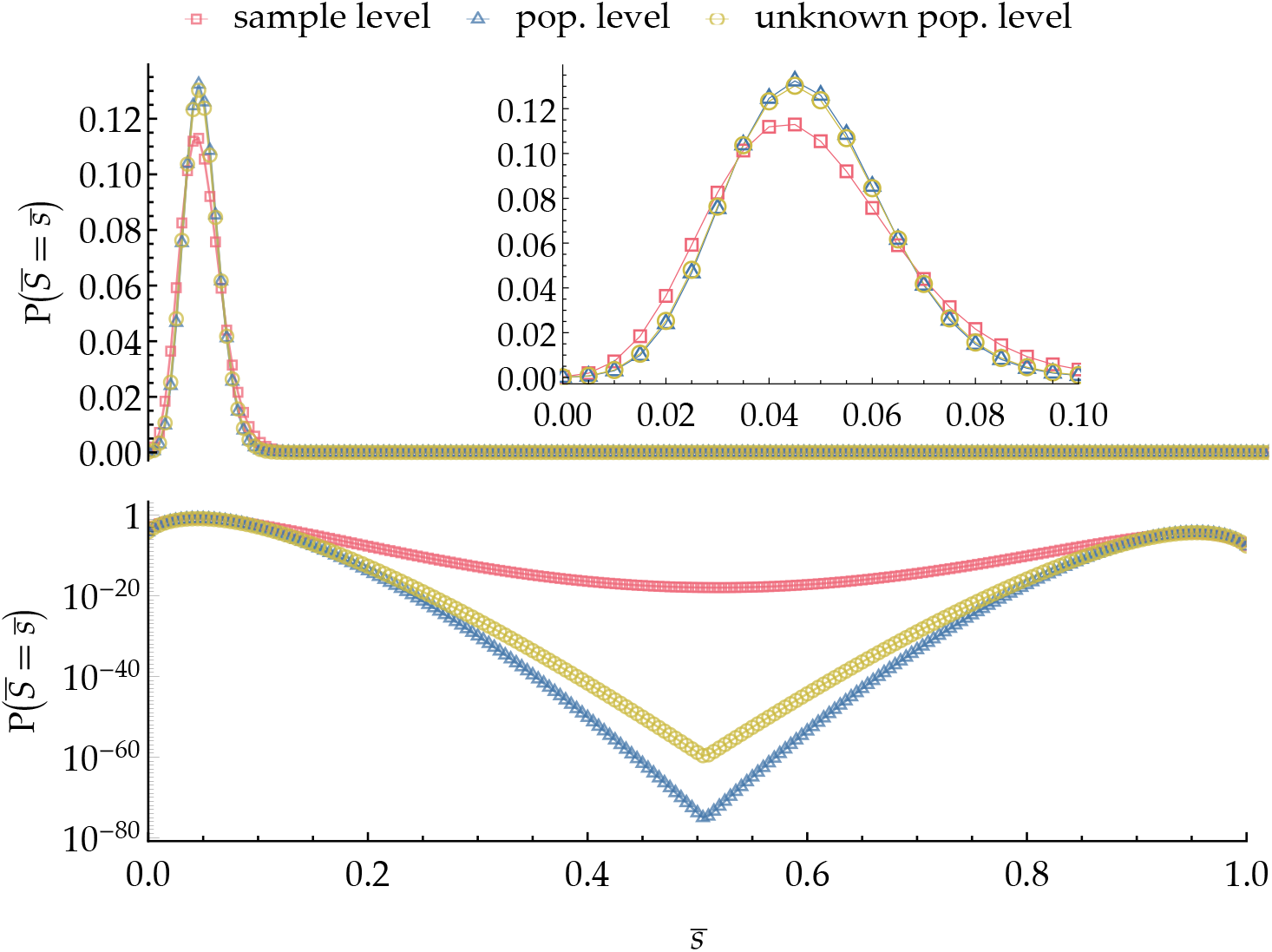
Linear and log-plots of 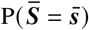 for a sample of *n* = 200 and constraints as in eq. (18), constructed by: **red squares:** maximum-entropy at the sample level, eq. (13); **blue triangles:** maximumentropy at the population level, eq. (17) with *ν* = 10 000, followed by sample marginalization; **yellow circles:** maximum-entropy at the population level with unknown population size, eq. (19), according to the distribution (20) for the population.

In our discussion we have so far assumed the size *ν* of the larger population to be known. This is rarely the case, however. We usually are uncertain about *ν* and can only guess its order of magnitude. In such a state of knowledge *I*_u_ our ignorance about the possible value of *ν* is expressed by a probability distribution P(*N* = *ν*|*I*_u_) = *h*(*ν*), and the marginal distribution for the sample activity (17) is modified, by the law of total probability, to

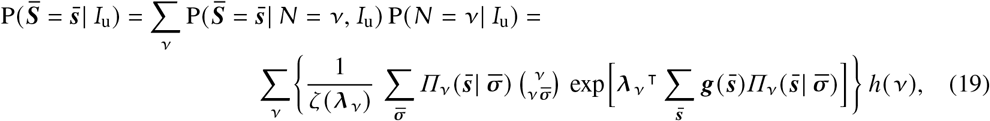

where the Lagrange multipliers ***λ***_*ν*_ and the summation range for 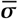 depend on *ν*.

As a proof of concept, fig. 2 also shows such a distribution (yellow circles) for the same constraints as above, and a probability distribution for *ν* inspired by Jeffreys [65 § 4.8]:

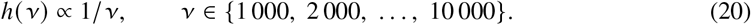

## 4 Discussion

The purpose of the present work was to point out and show, in a simple set-up, that the maximumentropy method can be applied to recorded neuronal data in a way that accounts for the larger population from which the data are sampled, eqs (15)–(17). This application leads to results that differ from the standard application which only considers the sample in isolation, eqs (13)–(14). We gave a numerical example of this difference. We have also shown how to extend the new application when the size of the larger population is unknown, eq. (19).

The latter formula, in particular, shows that the standard way of applying maximum-entropy implicitly assumes that *no* larger population exists beyond the recorded sample of neurons. One could in fact object to the application at the population level, and say that the traditional way of applying maximumentropy, eq. (13), yields different results because it does not make assumptions about the size *ν* of a possibly existing larger population. Such a state of uncertainty, however, is correctly formalized according to the laws of probability by introducing a probability distribution for *ν*, and is expressed by eq. (19). This expression cannot generally be equal to (13) unless the distribution for *ν* gives unit probability to *ν* = *n*; that is, unless the sample *is* the full population, and no larger population exists.

The standard maximum-entropy approach therefore assumes that the recorded neurons constitute a special subnetwork, isolated from the larger network of neurons in which it is embedded, and which was also present under the recording device. This assumption is unrealistic. The maximum-entropy approach at the population level does not make such assumption and is therefore preferable. It may reveal features in a data set that were unnoticed by the standard maximum-entropy approach.

The difference in the resulting distributions between the applications at the sample and at the population levels appears in the use of Boltzmann machines with hidden units [66], although by a different conceptual route. It also appears in statistical mechanics: if a system is statistically described by a maximum-entropy Gibbs state, its subsystems cannot be described by a Gibbs state [67]. A somewhat similar situation also appears in the statistical description of the final state of a non-equilibrium process starting and ending in two equilibrium states: we can describe our knowledge about the final state either by (1) a Gibbs distribution, calculated from the final equilibrium macrovariables, or (2) by the distribution obtained from the Liouville evolution of the Gibbs distribution assigned to the initial state. The two distributions differ (even though the final *physical* state is obviously exactly the same [68 § 4]), and the second allows us to make sharper predictions about the final physical state thanks to our knowledge of its preceding dynamics. In this example, though, both distributions are usually extremely sharp and practically lead to the same predictions. In neuroscientific applications, the difference in predictions of the sample vs full-population applications can instead be very relevant.

The idea of the new application leads in fact to more questions. For instance:

- Do the standard and new applications lead to different or contrasting conclusions about ‘cooperativity’, when applied to real data sets?
- How to extend the new application to the ‘inhomogeneous’ case [12; 13; 27], in which expectations for individual neurons or groups of neurons are constrained?
- What is the mathematical relation between the new application and maximum-entropy models with hidden neurons [36–39]?

Owing to space limitations we must leave a thorough investigation of these questions to future work.

Finally, we would like to point out the usefulness and importance of the probability formulae that relate our states of knowledge about a population and its samples, presented in § 2. This kind of formulae is essential in neuroscience, where we try to understand properties of extended brain regions from partial observations. The formulae presented here reflect a simple, symmetric state of ignorance. More work is needed [cf. 69] to extend these formulae to account for finer knowledge of the cerebral cortex and its network properties.

## Acknowledgements

PGLPM thanks Mari & Miri for continuous encouragement and affection; Buster Keaton for filling life with awe and inspiration; the developers and maintainers of LATEX, Emacs, AUCTEX, Open Science Framework, Python, Inkscape, Sci-Hub for making a free and unfiltered scientific exchange possible.

## Footnotes

1 Note that the ISO standard [42 § 3.1.14] states ‘The notion of representative sample is fraught with controversy, with some survey practitioners rejecting the term altogether’. Here we intend this notion in the Meaning 2 of Kruskal & Mosteller [43] or Meanings 8 and 9 of [44].

